# TPMS technology to infer biomarkers of macular degeneration prognosis in *in silico* simulated prototype-patients under the study of heart failure treatment with sacubitril and valsartan

**DOI:** 10.1101/625889

**Authors:** Guillem Jorba, Joaquim Aguirre-Plans, Valentin Junet, Cristina Segú-Vergés, José Luis Ruiz, Albert Pujol, Narcis Fernandez-Fuentes, José Manuel Mas, Baldo Oliva

## Abstract

Unveiling the mechanism of action of a drug is key to understand the benefits and adverse reactions of drug(s) in an organism. However, in complex diseases such as heart diseases there is not a unique mechanism of action but a wide range of different responses depending on the patient. Exploring this collection of mechanisms is one of the clues for a future personalised medicine. The Therapeutic Performance Mapping System (TPMS) is a Systems Biology approach that generates multiple models of the mechanism of action of a drug. This is achieved by (1) modelling the responses in human with an accurate description of the protein networks and (2) applying a Multilayer Perceptron-like and sampling method strategy to find all plausible solutions. In the present study, TPMS is applied to explore the diversity of mechanisms of action of the drug combination sacubitril/valsartan. We use TPMS to generate a range of mechanism of action models explaining the relationship between sacubitril/valsartan and heart failure (the indication), as well as evaluating their relationship with macular degeneration (a common/recurrent adverse effect). We found that a lower response in terms of heart failure treatment is more associated to macular degeneration development, although good response mechanisms can also associate to the adverse effect. A set of 30 potential biomarkers are proposed to identify mechanisms (or patients) more prone to suffering macular degeneration when presenting good heart failure response. As each molecular mechanism can be particular not only of cells but also individuals, we conclude that the study of the collection of models generated using TPMS technology can be used to detect adverse effects personalized to patients.

## Introduction

Cardiovascular diseases are the major cause of death in the Western world, causing 17.9 million deaths per year (2015).^1^ The prevalence of cardiovascular diseases is influenced by many factors: age, nutritional habits, lifestyles or genetics. This complicates the co-development of treatments and the identification of universal biomarkers to stratify the population. To facilitate this segmentation, it is necessary to understand the molecular details of the treatment and the pathology.

Sacubitril/valsartan (marketed by Novartis as Entresto®) is a drug combination that improves the results of the conventional treatments by reducing cardiovascular deaths and heart failure (HF) readmissions.^2^ In pharmacological terms, it is an angiotensin receptor-neprilysin inhibitor: it increases the natriuretic peptide system by inhibiting neprilysin (NEP) and inhibits reninangiotensin-aldosterone system by blocking the type-1 angiotensin II receptor (AT1R).^3^ In a previous work, the Mechanism of Action (MoA) of sacubitril/valsartan synergy was unveiled: the synergistic effect of the drug combination was mainly reflected on left ventricular extracellular matrix remodelling, mediated by proteins like gap junction alpha-1 protein or matrix metalloproteinase-9, and also affects cardiomyocyte apoptosis through modulation of glycogen synthase kinase-3 beta.^4^ However, several publications warned about the potential long-term negative implications of using a neprilysin inhibitor like sacubitril.^3,5,6^ Neprilysin plays a critical role at maintaining the amyloid-*β* homeostasis in the brain. The alteration of amyloid-*β* levels could lead to the development of Alzheimer’s disease or macular degeneration (MD).^7,8^ During the clinical trial PARADIGM-HF of sacubitril/valsartan, there were no serious effects detected.^2^ However, the patient follow-up of PARADIGM-HF was relatively short and not specialised in finding neurodegenerative specific symptoms. For this reason, in the recent sacubitril/valsartan clinical trial PARAGON-HF, there will be a Mini Mental State exam, and in the forthcoming PERSPECTIVE trial there will be a battery of cognitive tests^6^; however, any of the future studies will focus on potential MD consequences. A solution to explore the potential impact of sacubitril/valsartan on MD would be to apply computational methods that predict all the possible responses of the drug in the population. These computational simulations of real clinical trials are called *In Silico* Clinical Trials (ISCT), and they are based on systems biology principles.

Systems biology-based methods are increasingly becoming a reliable strategy to understand the molecular effects of a drug in complex clinical settings. However, current methodologies do not consider the inter-patient variability intrinsic to pharmacological treatments and thus miss relevant information that should be incorporated into the models. Indeed, there are many parameters influencing the MoA of such therapies, including demographic data of the patient, co-treatments or clinical history. The unexpected responses a patient might experience during a specific treatment could be explained by modeling the different mechanisms by which a drug exerts its effect on the patient. In this study, we will use the Therapeutic Performance Mapping System (TPMS)^9^ to elucidate all the possible MoAs of a sacubitril/valsartan in MD. TPMS is a systems biology approach based on the simulation of patient-like characteristics. It has been broadly used in the last years in different clinical areas and with different objectives.^4,10–14^ The TPMS incorporates data from massive databases publicly available and experimental information obtained for the diseases and drugs under study to generate multiple models of potential MoAs. Our hypothesis is that the set of MoAs could represent the different responses of the drug in cells and assume that a real population of patients is the result of a myriad of cell responses. Thus, a prototype-patient is defined as an abstract case with all cells responding with a single MoA. In this study, we analyse the population of MoAs associated with the response of HF and MD phenotypes to sacubitril/valsartan. We cluster the MoAs in groups by their response intensity. We analyse the MoAs with higher and lower susceptibility to treat HF and/or produce MD. We compare these MoAs and propose biomarkers to identify potential cases of MD when using sacubitril/valsartan. Finally, we use GUILDify v2.0 web server^15^ to analyse the comorbidity of both phenotypes and analyse the relevance of the proposed biomarkers with a different approach.

## Materials and Methods

### 1. Biological Effectors Database (BED) to molecularly describe specific clinical conditions

Biological Effectors Database (BED) describes more than 300 clinical phenotypes by means of gene and protein networks, which can be “active”, “inactive” or “neutral”.^10,16^ For example, in a metabolic network, proenzymes are “inactive” enzymes that become “active”, or enzymes are inactivated when they interact with an inhibitor (see further details in supplementary material).

### 2. TPMS modelling of phenotypes

The Therapeutic Performance Mapping System (TPMS) is a tool that creates mathematical models of a drug/pathology response to explain a clinical outcome or phenotype^4,9–14^. We applied TPMS to the drug-indication pair sacubitril/valsartan and HF: for the drug we retrieved and manually curated the sacubitril/valsartan targets from DrugBank^17^, PubChem^18^, STITCH^19^, SuperTarget^20^ and hand curated literature revision, while for the indication we used the proteins associated with the phenotype from the BED.^10,16^ We defined the Human Protein Network (HPN) by integrating data on functions and interactions of proteins (see further details in supplementary material).

#### 2.1 Defining restrictions from gene expression data

We define the state of human proteins as active or inactive for a particular phenotype, including its expression (as active) or repression (as inactive) extracted from the GSE57345 gene expression dataset^21^ as in *Iborra-Egea et al*^4^ (see further details in supplementary material).

#### 2.2 Description of the mathematical models

The algorithm of TPMS uses as input signals the values of activation (+1) and inactivation (−1) of the drug protein targets. The outputs are then the values of activation and inactivation of the proteins defining the indication’s phenotype (retrieved from the BED), named effectors. Each node of the protein network receives as input the output of the connected nodes and each link receives a weight (ω_*l*_). The sum of inputs is transformed by a hyperbolic tangent function to generate the score of the node, which becomes the “output signal” towards the connected nodes. The ω_*l*_ parameters are obtained by optimization, using a Stochastic Optimization Method based on Simulated Annealing.^22^ The models are trained by using the restrictions defined by the BED and the specific conditions set by the user. We ranked all solutions by the number of restrictions satisfied and selected the top 200 solutions representing potential MoAs of the drug, which we assumed equally acceptable. Here, we hypothesized that these solutions can represent different cells, while combinations of them would correspond to different patients. Hence, 200 prototype or representative mathematical solutions can be considered for an individual and personalized approach. Details of the approach are shown in **Figure 1** and supplementary material.

**Figure 1.**
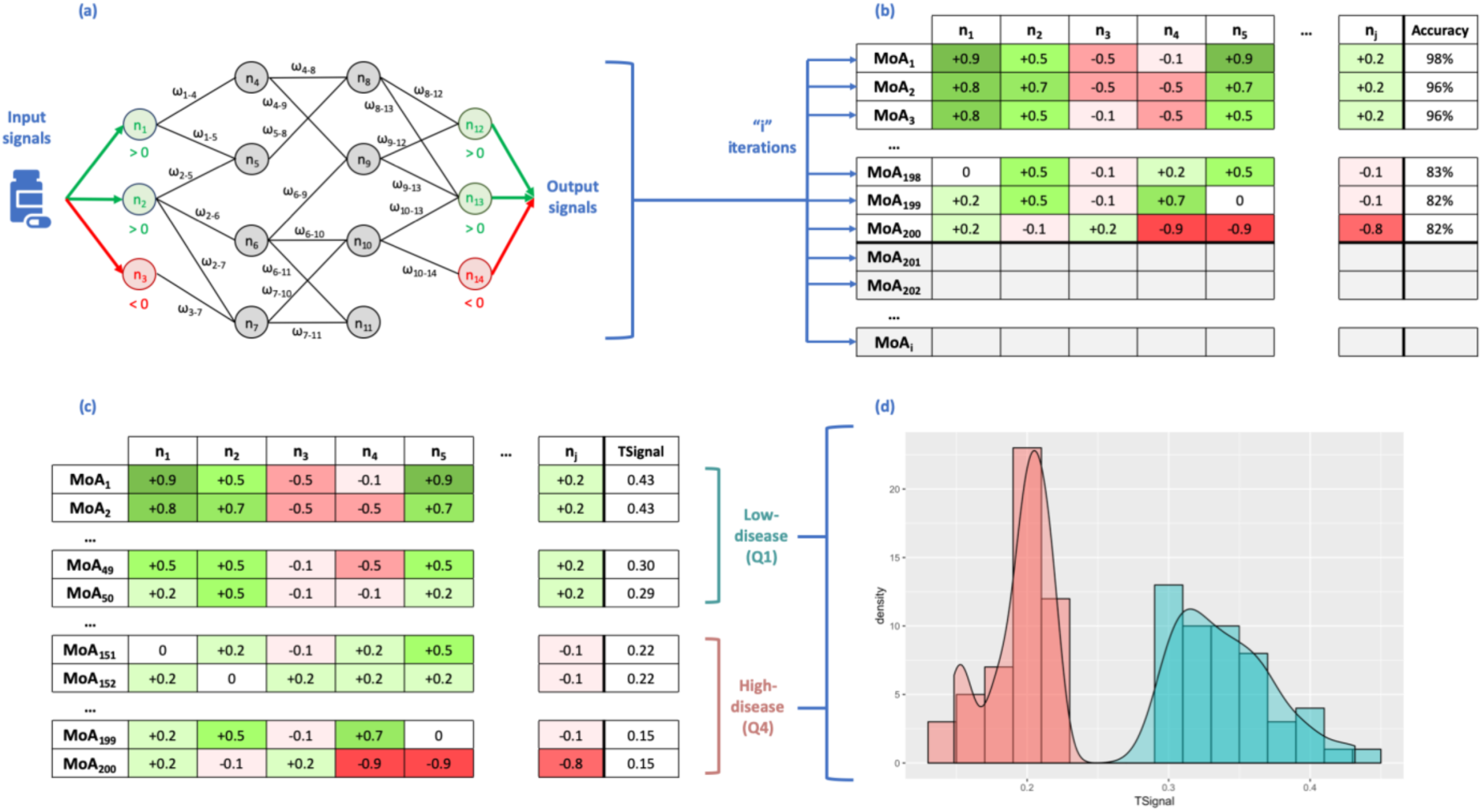
Scheme of the method, transmitting information over the Human Protein Network (HPN) using a Multilayer Perceptron-like and sampling **(a)**. After a given number of iterations, we obtain a collection of Mechanisms of Actions (MoA) **(b)**. Rows represent the MoAs and columns the output signal values of the proteins (nodes of the network). The final column shows the accuracy of the model as a percentage of the number restrictions accomplished and 200 MoAs are selected (coloured in the slide) and sorted by TSignal. The first quartile is defined as the Low-disease group, and the fourth quartile as High-disease group **(c)**. The distribution of the output signals of the two groups of MoA are shown in **(d)** (High-disease in red and Low-disease is in blue).

### 3. Measures to compare sets of MoAs

To understand the relationships between all potential mechanisms we defined measures of comparison between different sets of solutions. We expect that a drug will revert the conditions of a disease phenotype. Consequently, a drug should inactivate the active protein effectors of a pathology-phenotype and activate the inactive ones. Here, we defined several measures in order to study and compare sets of MoAs from different views (see further details in supplementary material).

#### 3.1 TSignal

To quantify the intensity of the response of a MoA and compare it with others, we created a measure called TSignal. The TSignal is the average of the output signals of the protein effectors (equation in supplementary material).

#### 3.2 Distance between two sets of MoAs

We calculated the *distance* between two or more sets of MoAs in order to determine their similarity. For that, we used a modified Hausdorff distance (MHD) introduced by Dubuisson and Jain.^23^ Details of equations are in supplementary material.

#### 3.3 Potential biomarkers extracted from MoAs

For HF, MoAs are ranked by their TSignal and split in four quartiles: the first quartile (top 25%) contains MoAs with higher intensity of the response (TSignal), which in turn reduces the values of the effectors associated with a disease phenotype (we named them as Low-disease MoAs). The fourth quartile (bottom 25%) collects MoAs with lower intensity of response (thus, we named as High-disease MoAs). On the other hand, for MD, the first quartile (top 25%) contains MoAs with higher intensity, which in this case, as an adverse event, it increases the values of the effectors associated to the comorbidity (we name them as High-adverseEvent MoAs). The fourth quartile (bottom 25%) collects MoAs with lower intensity of response (thus, we named as Low-adverseEvent MoAs).

We use the comparison between both groups of High-and Low-MoAs to identify the *best-classifier proteins*. Best-classifier proteins are those that classify the best between High-and Low-groups and are determined by a Data-Science strategy (see supplementary material). Best-classifier proteins are strongly related to the intensity of a response and are differently distributed between High-and Low-MoAs. We only select the 200 proteins (or pair of proteins) reproducing with higher accuracy the classification. Assuming the hypothesis that the selected MoAs are representative of individual prototype patients, these proteins can be used as biomarkers to classify a cohort of patients by the activity or inhibition of the protein.

Each best-classifier protein has different output signals in the Low-and High-group MoAs and the distribution in both sets can be compared. We use a Mann-Whitney *U* test to compare the two distributions of output signals and select those proteins for which the difference is significant (p-value< 0.01), having an average output signal in Low-HF with opposite sign to the average output signal in High-HF (i.e. positive vs. negative or vice versa). We name these as *differential best-classifier proteins*. By following this strategy, we can identify two groups of differential best-classifier proteins: those active in Low-disease (positive output signal in average) and inactive in High-disease (negative output signal in average), and those active in High-disease and inactive in Low-disease.

## Results and discussion

We applied TPMS to the HPN using as input signals the drug targets of sacubitril/valsartan (NEP / AT1R) and as output signals the proteins associated with Heart Failure (HF) extracted from the BED. As described in the methodology, out of all MoAs found by TPMS, we selected 200 satisfying the largest number of restrictions (and at least 80% of them) to perform further analysis. To rank the MoAs according to the intensity of the signal arriving from the drug, we calculated the TSignal of every MoA associated with HF and MD, i.e. the average output signal arriving to the protein effectors of both pathologies. According to the TSignal of HF and following the procedure described in Materials and Methods, we defined two groups of MoAs: *Low-HF*, containing the MoAs with a higher intensity of the response and therefore a healthier phenotype, and *High-HF*, with the MoAs of lower intensity of the response and therefore an increased HF disease phenotype. We also calculated the TSignal of MD and define two groups of MoAs: *Low-MD*, producing a reduced adverse effect, and *High-MD*, producing an increased adverse effect.

Note that TPMS was only executed once, optimising the results to satisfy the restrictions on HF data. The values of MD are obtained by measuring the signal arriving at the MD effectors, which are part of the HPN and also receive signal. This procedure was chosen because we treat HF as the indication of the drug (sacubitril/valsartan), while MD is a potential adverse effect.

In the following sections, we analysed and compared the four groups of MoAs and searched for biomarkers that can potentially identify MD as an adverse effect of the drug (and consequently classify patients). We used the web server GUILDify v2.0^15^ to analyse the comorbidity of both phenotypes and analyse the relevance of the proposed biomarkers in a different context.

### 1. Analysis of MoAs of high/low intensity response associated to HF

We ranked the MoAs by the TSignal on HF effectors. Consequently, the MoAs on the set High-HF corresponds to those with lower signal affecting the effectors of HF, while MoAs on the set Low-HF are the opposite. Most of the High-HF models have TSignal between 0.15 and 0.25, while for Low-HF TSignal ranges between 0.3 and 0.4 (**Supplementary Figure 1a**).

We selected the 200 best-classifier proteins after defining the two groups of MoAs. These are defined as the proteins (or pairs of proteins) that can allocate better the MoAs in High-HF and Low-HF. Among these proteins, we identified the differential best-classifier proteins. These are proteins that have output signals significantly different between Low-HF and High-HF (Mann-Whitney *U* test, adjusted p-value<0.01) and for which the average has opposite sign between the cohorts (i.e. active in Low and inactive in High or vice versa). We identified two groups of differential best-classifier proteins: those active in Low-HF (the average of output signals in Low-HF MoAs is positive) and inactive in High-HF (the average of output signals in High-HF MoAs is negative), and those active in High-HF and inactive in Low-HF. Out of the starting 200 best-classifier proteins, 45 are differential (6 in the first group and 39 in the second) (see **Supplementary Table 1**). **Figure 2a** shows a plot for all the proteins in the MoAs and their average output signals in the MoAs of Low-HF and High-HF. Most of the proteins with opposite signs between the cohorts are found as best-classifier proteins.

We calculated the Gene Ontology (GO) enriched functions of the differential best-classifier proteins that are active in Low-HF MoAs and inactive in High-HF MoAs. The enrichment was calculated using the software FuncAssociate.^24^ Among the enriched functions we find processes associated with the SCAR complex, the positive regulation of actin nucleation, the regulation of neurotrophin TRK receptor and dendrite extension. We did the same for GO functions of the differential best-classifier proteins that are inactive in Low-HF and active in High-HF. We find functions such as phosphatidylinositol kinase activity, MAP kinase activity, DNA damage induced protein phosphorylation and superoxide anion generation. Although some functions are enriched in both sets, such as Fc gamma receptor signalling, the majority of functions enriched in different groups of differential best-classifier proteins are different (see **Supplementary Table 2)**.

Some of the proteins and functions highlighted in the current analysis have been related to myocardial function. On the one hand, our differential best-classifier proteins active in Low-HF MoAs and inactive in High-HF MoAs point towards an important role for actin nucleation and polymerization mechanisms in drug response (reflected by the functions *regulation of actin nucleation, regulation of Arp2/3 complex-mediated actin nucleation, SCAR complex, filopodium tip*, or *dendrite extension*); in fact, the alteration of actin nucleation and polymerization mechanisms has been reported in heart failure.^25–27^ Interestingly, a role for the activation of another differential best-classifier candidate, ATGR2, has been proposed to mediate some of the beneficial effects of angiotensin II receptor type 1 antagonists, such as valsartan.^28,29^

On the other hand, results for the differential best-classifier proteins inactive in Low-HF and active in High-HF point towards the importance of phosphatidylinositol kinase mediated pathways (*phosphatidylinositol-3,4-bisphosphate 5-kinase activity*) and MAP kinase mediated pathways (*MAP kinase kinase activity*, best classifier proteins MAPK1, MAPK3, MAPK11, MAPK12 or MAPK13) in response to sacubitril/valsartan; furthermore, both signalling pathways have been associated to cardiac hypertrophy and subsequent heart failure.^30,31^

### 2. Analysis of MoAs of high/low TSignal associated to MD

We also ranked the MoAs by the TSignal on MD effectors. We similarly classified MoAs in High-MD and Low-MD. High-MD involve MoAs with high signal and higher probability to develop the adverse effect phenotype, while MoAs on the set Low-MD corresponds to healthier MoAs. TSignal ranges between 0.20 and 0.30 for Low-MD models and between 0.35 and 0.45 for High-MD (**Supplementary Figure 1b**).

We identified the 200 best-classifier proteins to divide MoAs in Low-MD and High-MD. We also identified the differential best-classifier proteins active in Low-MD and inactive in High-MD and vice versa. As before, we compared the distributions of Low-MD and High-MD output signals of the best-classifier proteins and calculate the average of the signal in all MoAs in Low-and High-MD. Out of 200 best-classifier proteins, we identified 28 active in Low-MD (inactive in High-MD) and 29 active in High-MD (inactive in Low-MD) (see **Supplementary Table 3**). **Figure 2b** shows the plot for all proteins classified by their average output signal in Low-MD and High-MD models.

**Figure 2.**
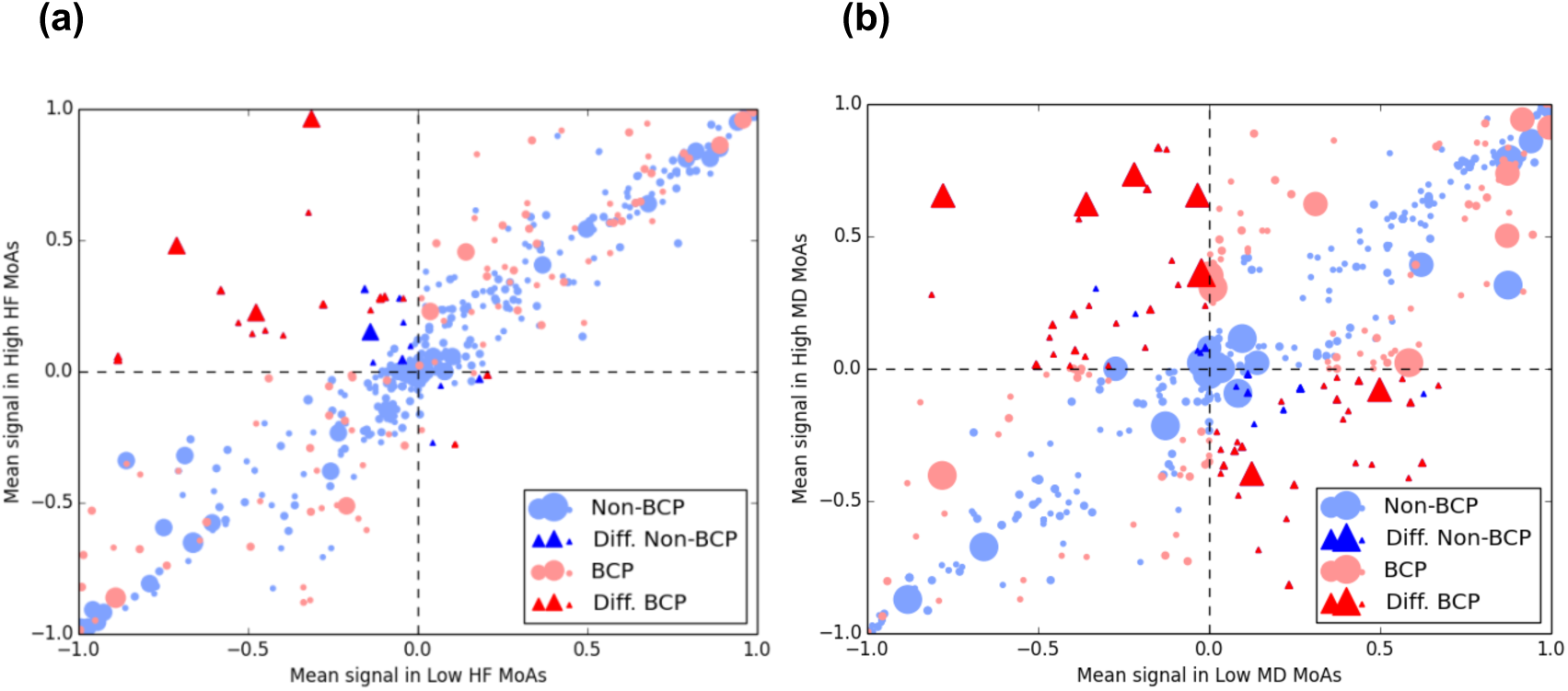
Scatter plot of the mean signal values of Low-“disease” and High-“disease” MoAs for each protein using as disease Heart Failure (HF) in **(a)** and Macular Degeneration (MD) in **(b)**. The average of the output signal of each protein in High-group is presented versus its value in Low-group. Differential signals (Diff., shown as triangles) are defined as those with opposite sign when comparing High versus Low average, and a p-value < 0.01 when calculating the Mann-Whitney *U* test between the two distributions of signals. Best-classifier proteins (BCP) are coloured in red, otherwise they are blue. Sizes of markers are proportional to GUILDify score.

We calculated the GO enriched functions for these groups of proteins (see **Supplementary Table 4**). For the first group (Low-MD active / High-MD inactive) we obtain unique functions such as dendritic spine development, positive regulation of vascular endothelial growth factor production and phosphotyrosine binding. For the second group (Low-MD inactive / High-MD active) we observe functions such as dorsal/ventral axon guidance, fibroblast growth factor receptor binding and response to toxic substance. Then, phosphatidylinositol bisphosphate kinase activity is clearly the enriched function in both groups.

Some of the proteins and functions highlighted in the current analysis have been related to MD. The presence of dendritic spine development related proteins among the differential best-classifiers proteins (Low-MD active and High-MD inactive) and dorsal/ventral axon guidance related proteins among the inverse classifiers (Low-MD inactive and High-MD active) points towards a role for sacubitril/valsartan-associated MD in dendritic and synaptic plasticity mechanisms, which have been previously linked to the condition.^32^ Furthermore, valsartan treatment has been reported to promote dendritic spine development in other related neurodegenerative diseases, such as Alzheimer’s disease.^33^ Other functions enriched within the differential best-classifier proteins (Low-MD inactive and High-MD active) are implicated in growth factor related pathways, which are known to be involved in wet MD pathogenesis.^34^ Moreover, neovascularization in the wet variant of MD has been linked to the signalling of some of the growth factors detected as sacubitril/valsartan-associated MD classifiers in this study, including FGF1^34^ and PDGF^35,36^.

### 3. Comparison of MoAs of high/low TSignal associated with HF and MD

We calculated the modified Hausdorff distance between the sets of MoAs in High-MD, Low-MD, High-HF and Low-HF (**Supplementary Table 5**). It is remarkable that the distances between Low-HF and High-HF and between Low-MD and High-MD are larger than the cross distances between HF and MD. We used these distances to calculate a dendrogram tree (see **Supplementary Figure 2**) showing that MoAs associated with a bad response of sacubitril/valsartan for HF (high-HF) are more similar (i.e. closer) to MoAs linked to a stronger MD adverse effect (high-MD) than those corresponding to Low HF and MD. However, by the definition of distance (equation 3 in supplementary material), we cannot account for the dispersion among the MoAs within and between each group. Therefore, we calculated for each set the mean Euclidean distance between all the points and its centre defined by the average of all (see **Supplementary Table 6**). We expect to find common MoAs between different HF and MD-defined sets, meaning that some MoAs belonging to one of the two HF sets are also found in one of the MD sets.

In order to have a global view of the distance between the MoAs in sets High-and Low-of HF and MD, we used the multidimensional scaling (MDS) plot calculated with the MATLAB function “mdscale” and the metric scaling “metricsstress” (see **Figure 3**). MDS plots pairwise distances in two dimensions that preserve the clustering characteristics (i.e. close MoAs are also close in the 2D-plot and far MoAs are also far in 2D). Focusing on Low-HF, depicted in orange, we observe that, despite its dispersion, some MoAs are close to High-MD, while others are closer to the Low-MD set. This implies that the response to sacubitril/valsartan of HF patients, in the best scenario when the TSignal of the disease effectors is decreased, it does not necessarily imply an improvement on the intensity of MD (i.e. Low-MD). Assuming the hypothesis that different MoAs correspond with prototype patients, we can conclude that taking sacubitril/valsartan could produce MD in a scarce number of patients. When focusing in the set of High-HF MoAs, we differentiate between two clusters of MoAs: one related to the High-MD set; and other MoAs close to MoAs in the “Low-MD” set. Assuming that each MoA corresponds with a prototype patient, we conclude that for a specific set of patients sacubitril/valsartan does not reduce HF and increases the MD TSignal while for others it does not produce MD. Such view of the MoAs of sacubitril/valsartan can help us stratify patients depending on the biomarkers that characterize the sets of MoAs as seen in previous sections.

**Figure 3.**
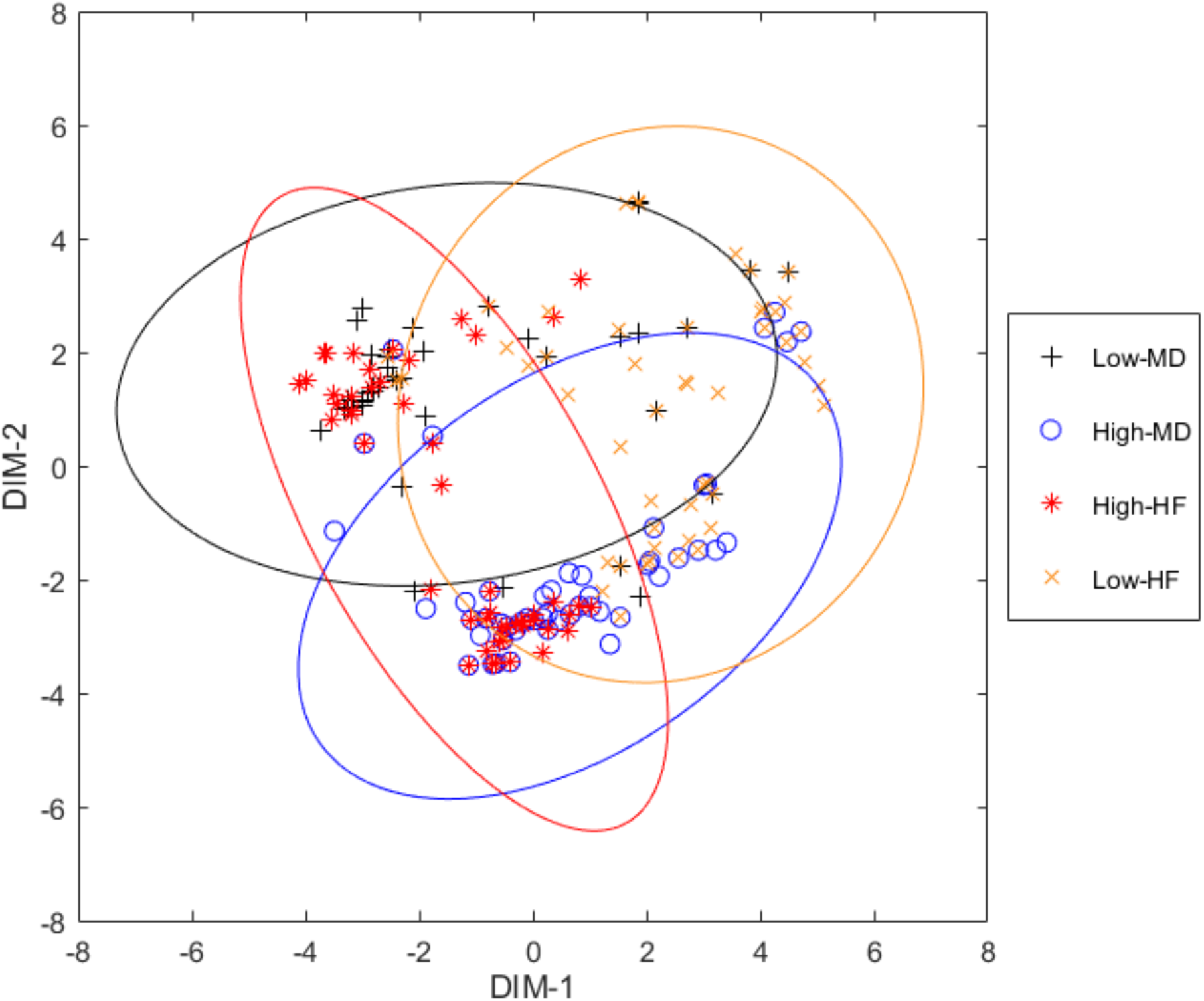
Multidimensional scaling plot of the distances between the MoAs of the four groups defined. Each point represents a MoA. Axes are defined by the most representative dimensions.

We calculated the number of common MoAs for High-and Low-HF and High-and Low-MD (**Supplementary Table 7**). High-HF and High-MD have the largest number of common MoAs (17), while Low-HF and High-MD have the lowest (9). Focusing on Low-HF, we analysed those proteins potentially responsible of MD adverse effect (i.e. High-MD). We analysed the distribution of output signals of proteins in common MoAs of Low-HF and Low-MD (Low-HF ∩Low-MD) and the distribution of output signals of proteins in common MoAs of Low-HF and High-MD (Low-HF ∩ High-MD). We selected those nodes (i.e. proteins) with significant differences (using a Mann-Whitney *U* test) for which the averages of output signals have opposite signs in both sets (see methods in 3.3). **Table 1** shows 30 proteins selected, which are the potential biomarkers to identify MoAs of High-MD. On the one hand, 16 of them are active in Low-HF ∩ Low-MD MoAs and inactive in Low-HF ∩ High-MD. We note that 15 of these are best-classifier proteins of MD. On the other hand, 14 proteins are inactive in Low-HF ∩ Low-MD and active in Low-HF ∩ High-MD MoAs, and 12 of them are best-classifier proteins of MD. We calculated the GO enriched functions of these two groups, observing that “phosphatidylinositol bisphosphate kinase activity” is the function enriched among proteins that are active in Low-HF ∩ Low-MD MoAs, while “fibrinolysis” is the enriched function among proteins active in Low-HF ∩ High-MD MoAs (**Table 2**). With this, we conclude that while trying to improve the health of patients for HF, in specific patients we may induce MD through the modulation of fibrinolysis (specifically involving 12 best-classifier proteins that may be considered as good biomarkers for this prognosis).

**Table 1.**
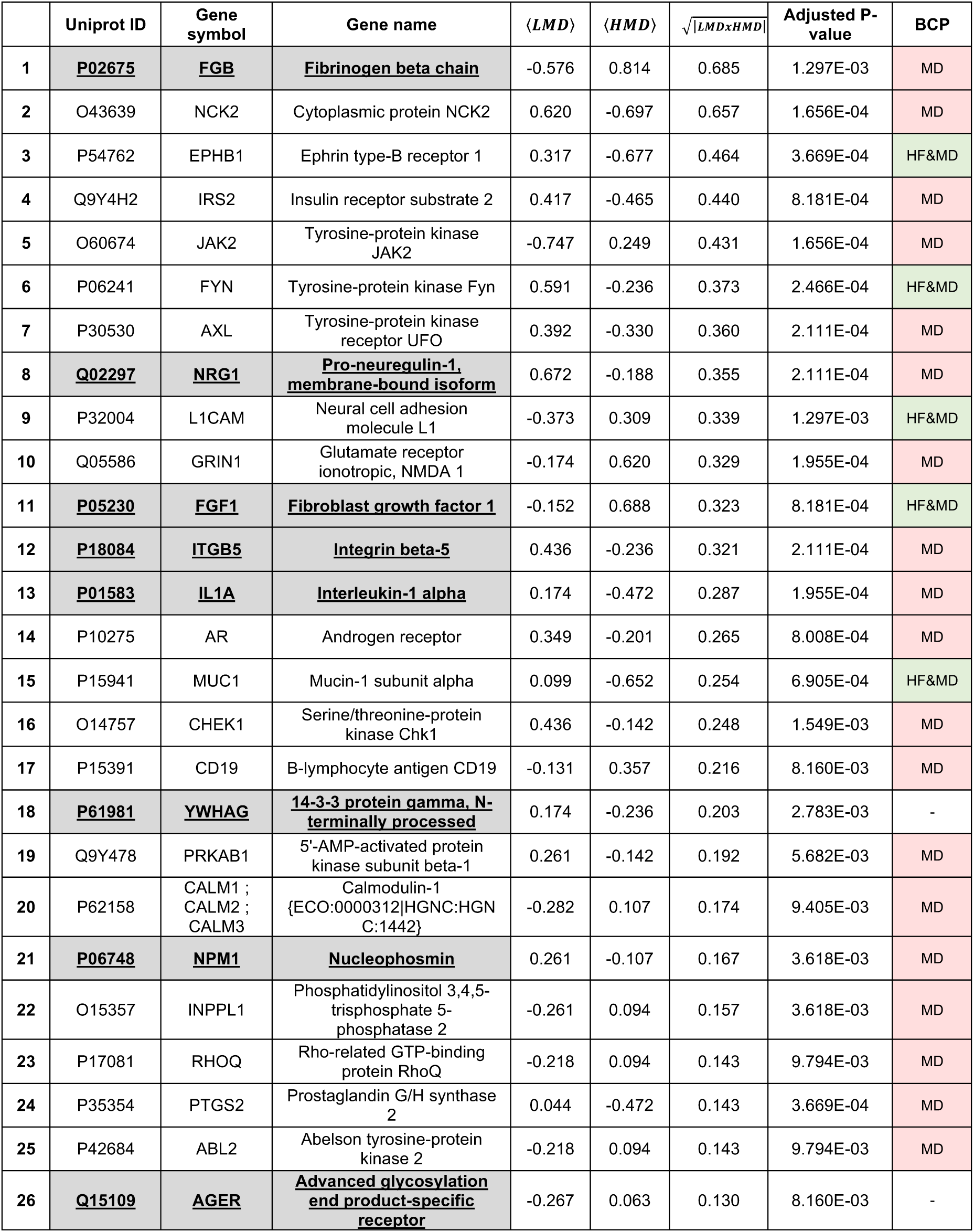
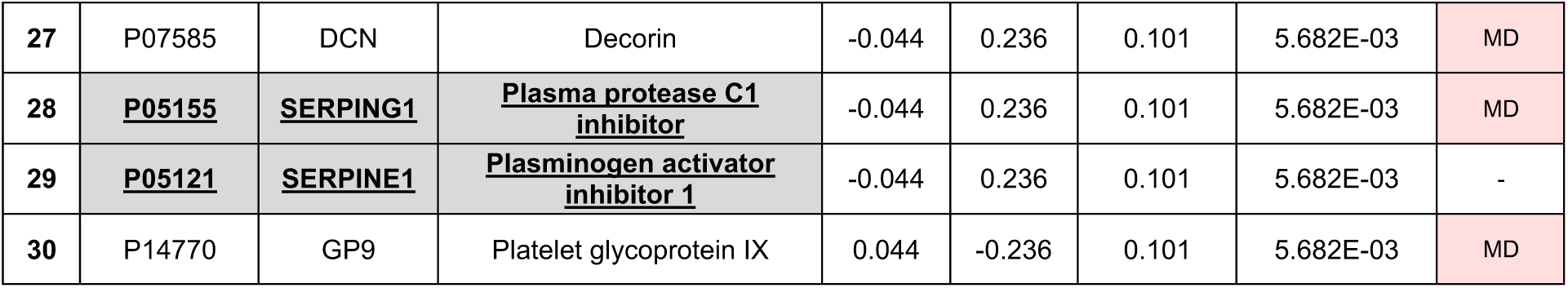
Biomarker proteins, with opposite signal in Low-HF ∩ Low-MD and Low-HF ∩ High-MD MoAs. Highlighted cells correspond to proteins that are part of the Top-HF υTop-MD υTop-Drug set, the top-scoring proteins according to GUILDify. Columns show: the protein name (as UniprotID, gene-symbol and gene-name), the average of the signal in in Low-MD (<LMD>) and High-MD (<HMD>) in the selected sets of MoAs and a measure of the strength of the signal in both distributions (calculated as *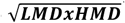*), the significance (adjusted P-value) ensuring that both distributions of signals are different, and whether the protein has been considered best-classifier in MD of HF (BCP).

**Table 2.** Top 10 gene Ontology functions enriched from proteins with opposite signal in Low-HF ∩ Low-MD and Low-HF ∩ High-MD MoAs. Functional enrichment analysis from FuncAssociate[REF].

| | Low-HF ∩ LMD+ HMD- | | | Low-HF ∩ HMD+ LMD- | | | Overlapped functions | | |
| --- | --- | --- | --- | --- | --- | --- | --- | --- | --- |
|  | GO name | LOD | P-val. | GO name | LOD | P-val. | GO name | LOD | P-val. |
| 1 | phosphatidylinositol-4,5-bisphosphate 3-kinase activity | 1.89 | 0.03600 | fibrinolysis | 2.51 | 0.00050 | response to stimulus | 1.19 | <0.00050 |
| 2 | cellular response to UV | 1.87 | 0.04200 | negative regulation of wound healing | 2.13 | 0.00050 | positive regulation of transport | 1.24 | <0.00050 |
| 3 | phosphatidylinositol bisphosphate kinase activity | 1.87 | 0.04200 | negative regulation of blood coagulation | 2.12 | 0.00850 | positive regulation of biological process | 1.13 | 0.00051 |
| 4 | vascular endothelial growth factor receptor signaling pathway | 1.86 | 0.04200 | negative regulation of hemostasis | 2.12 | 0.00850 | positive regulation of developmental process | 1.18 | <0.00050 |
| 5 | positive regulation of protein kinase B signaling | 1.70 | 0.01050 | negative regulation of coagulation | 2.10 | 0.01050 | positive regulation of cellular process | 1.04 | 0.00294 |
| 6 | negative regulation of apoptotic signaling pathway | 1.68 | 0.00050 | platelet alpha granule lumen | 1.96 | 0.02300 | positive regulation of response to stimulus | 1.04 | 0.00417 |
| 7 | peptidyl-tyrosine phosphorylation | 1.63 | 0.01400 | regulation of epithelial cell apoptotic process | 1.96 | 0.02300 | - | - | - |
| 8 | regulation of apoptotic signaling pathway | 1.63 | <0.00050 | regulation of blood coagulation | 1.91 | 0.02800 | - | - | - |
| 9 | peptidyl-tyrosine modification | 1.62 | 0.01400 | regulation of hemostasis | 1.91 | 0.02800 | - | - | - |
| 10 | protein tyrosine kinase activity | 1.61 | 0.01850 | regulation of coagulation | 1.89 | 0.03450 | - | - | - |

In fact, since neovascular MD development is characterized by subretinal extravasations of novel vessels derived from the choroid (CNV) and the subsequent haemorrhage into the photoreceptor cell layer in the macula region^37^, it might be reasonable to think that the modulation of fibrinolysis and blood coagulation pathways might play a role. The reported implication of some fibrinolysis related classifiers, such as FGB, SERPINE1 (PAI-1), and SERPING1, in neovascular MD development seems to support this hypothesis.^38–40^ Moreover, valsartan might be implicated in this mechanism, since it has been reported to modulate PAI-1 levels and promote fibrinolysis in different animal and human models.^41,42^

In addition, the presence of several other MD related classifiers in this list, such as IRS2^43^, PTGS2^44^, DCN^45^ and FGF1^46^, further supports the interest of the classifiers as biomarkers of MD development in sacubitril/valsartan good responders.

### 4. Analysis of comorbidity and potential biomarkers with GUILDify

We used GUILDify v2.0^15^ to deep on the potential comorbidity between HF and MD. We proved in previous works the use of gene-prioritization algorithms to study the mechanisms involved in comorbiditites.^47^ The principle of the approach is that GUILDify v2.0 extends the information of disease-gene associations through the PPI network and helps to connect the original associations (seeds) through the network. As seeds, we used the effectors of the BED database: 124 seeds for HF and 163 seeds for MD. We selected the top 2% nodes scored by GUILDify (i.e. 262 nodes for each patho-phenotype, named “top-HF” and “top-MD” respectively). We obtained 36 proteins and 19 GO enriched biological processes in common, suggesting the potential relationship of both diseases (see more details in **Supplementary Figure 3**).

The top scoring nodes selected by GUILDify tend to have an important role on transmitting the information of the phenotype through the network. Thus, we used GUILDify to indicate which of the differential best-classifier proteins identified by TPMS may have a relevant role in the phenotype. We ran GUILDify using the two targets of sacubitril/valsartan (NEP, AT1R) as seeds, and selected the top 2% scored nodes (defined as the “top-drug” set). We merged the top scored sets of HF, MD and top-drug (“top-drug ∪ top-HF ∪ top-MD”) and studied the overlap with the set of differential best-classifier proteins of the MoAs associated with MD and HF. **Supplementary Table 8** shows the result of this analysis, with a significant representation of best-classifier proteins in most of the sets, especially significant on MD best-classifier proteins. In **Figure 2**, the differential best-classifier proteins with higher score can be identified by a larger area. **Supplementary Table 9** shows the list of 13 proteins involved in this overlap. We have also checked the overlap with the 30 biomarkers proposed in the previous section, of which 10 are found in the merged set “top-drug ∪ top-HF ∪ top-MD” and are consequently significant (see **Supplementary Tables 10 and 11**).

Some of these candidates can be functionally linked to both diseases and the drug under study. For example, among these 10 classifiers, AGER has been implicated in both HF^48^, through extracellular matrix remodelling, and MD development^49^, through inflammation, oxidative stress, and basal laminar deposit formation between retinal pigment epithelium cells and the basal membrane; furthermore, this receptor is known to be modulated by AT1R^50^, valsartan target. Similarly, FGF1 has been proposed to improve cardiac function after HF^51^, as well as to promote choroid neovascularization leading to MD.^34^ Moreover, FGF1 is regulated by angiotensin II through ATGR2^52^, another protein suggested as classifier in the current analysis that is known to mediate some of the effects of AT1R antagonists, such as valsartan.^28,29^ Another candidate, NRG1, has been linked to myocardial regeneration after HF^53^ and is known to lessen the development of neurodegenerative diseases such as Alzheimer’s disease^54^, which shares similar pathological features with MD^55^. NRG1 is also linked to the expression of neprilysin^54^, sacubitril target. ITGB5 has been identified as risk locus for HF^56^ and its modulation has been linked to lipofucsin accumulation in MD^57^. Interestingly, ATGR1 inhibitors have been reported to modulate ITGB5 expression in animal models^58^. Finally, IL1A has been proposed as an essential mediator of HF pathogenesis^59,60^ through inflammation modulations, and serum levels of this protein have been found increased in MD patients.^61^ In addition, as described in previous sections, classifiers FGB, SERPINE1, and SERPING1 have been linked to MD^38–40^ and are also known to play a role in HF development.^62–65^

According to these findings, the identified classifiers might represent potential biomarkers for the identification of sacubitril/valsartan good responder HF patients at risk of MD development.

## Limitations

Although TPMS returns the amount of signal from the drug arriving to the rest of the proteins in the HPN, this signal is only a qualitative measure. We are not using data about the dosage of the drug or the quantity of expression of the proteins. However, we are already working to make TPMS move towards the growing tendency of Quantitative Systems Pharmacology. The quantification of the availability of drugs in the target tissue for each patient opens the opportunity to have an accurate patient simulation to do *in silico* clinical trials.

## Conclusions

There is a need of systems biology-based methods to simulate the different responses of a drug in patients. The specific case of sacubitril/valsartan stands out because of the amount of resources invested in the safety of the drug and the concern on the risk of MD. In this study, we apply TPMS to uncover the different MoAs of sacubitril/valsartan over HF and reveal its relationship with MD. We hypothesize that all MoAs coexist in cells, but in a population of patients some cells may have more proclivity to certain MoAs than others. We define a prototype-patient as one with a single MoA for a drug and study the *in silico* trial of HF treatment with sacubitril/valsartan and its potential adverse effect MD. TPMS achieves this by modelling an accurate representation of the HPN and applying a Multilayer Perceptron-like and sampling method strategy to find all plausible solutions. We found that HF low responder MoAs are more associated to MD development at the same time, although good responders are also associated to MD. Different sets of proteins have been found to classify the mechanisms according to HF and MD response, which include functions such as PI3K and MAPK kinase signalling pathways, involved in HF-related cardiac hypertrophy, or fibrinolysis and coagulation processes (e.g. FGB, SERPINE1 or SERPING1) and growth factors (e.g. FGF1 or PDGF) related to MD induction. We propose 30 biomarkers to identify patients potentially developing MD under the successful treatment with sacubitril/valsartan. Out of this 30, we propose 10 biomarkers that have been found to be involved in the comorbidity between HF and MD predicted by a different approach (GUILDify), including well-known HF and MD effectors also related to the mechanisms of sacubitril/valsartan and/or HF, such as AGER, NRG1, ITGB5 or IL1A.

## Authorship Confirmation Statement

JAP, GJ, CSV, JMM & BO analysed the data. GJ, VJ, CSV, JLR & JMM studied HF and MD with TPMS. AP, GJ, JLR & JMM described the methods of TPMS. JAP, NFF & BO described the methods of GUILDify. CSV & JMM analysed and described the potential biomarkers for MD adverse effects by sacubitril/valsartan. JAP, NFF & BO compared the results of TPMS with GUILDify. All authors were involved in writing the manuscript.

## Author Disclosure Statements

GJ, VJ, CSV, AP, JLR & JMM have financial interest on the commercial package of TPMS and the database of effectors (BED).

JAP, NFF & BO don’t have any competing financial interests.

## Abbreviations

TPMS: Therapeutic Performance Mapping System
HF: Heart Failure
MD: Macular Degeneration
MoA: Mechanism of Action
ISCT: *In Silico* Clinical Trials
BED: Biological Effectors Database
HPN: Human Protein Network
GO: Gene Ontology

## Acknowledgements

The authors received support from: the Spanish Ministry of Economy (MINECO) [BIO2017-85329-R] [RYC-2015-17519]; “Unidad de Excelencia María de Maeztu”, funded by the Spanish Ministry of Economy [ref: MDM-2014-0370]. The Research Programme on Biomedical Informatics (GRIB) is a member of the Spanish National Bioinformatics Institute (INB), PRB2-ISCIII and is supported by grant PT13/0001/0023, of the PE I+D+i 2013-2016, funded by ISCIII and FEDER. GJ and VJ have received funding from the European Union’s Horizon 2020 research and innovation programme under the Marie Sklodowska-Curie grant agreement [refs: 765912; 765158].

