## supplementary for "TPMS technology to infer biomarkers of macular degeneration prognosis in *in silico* simulated prototype-patients under the study of heart failure treatment with sacubitril and valsartan"

### Extended version of material and methods

#### 1. Biological Effectors Database (BED) to molecularly describe specific clinical conditions

Patient-like characteristics are modelled using clinical data and/or experimental molecular data. There are many databases providing clinical data of patients, adverse drug reactions, diseases or indications (e.g. ClinicalTrials.gov, SIDER, ChEMBL, PubChem, DrugBank…). Many other databases provide molecular data, defining the existing human genes and/or proteins and describing the relationships between them (IntAct, BioGRID, REACTOME…). Combining both, the clinical and the molecular information available, the BED describes more than 300 clinical phenotypes by means of gene and protein networks, which can be "active", "inactive" or "neutral".[1,2] For example, in a metabolic network, proenzymes are "inactive" enzymes that become "active", or enzymes are inactivated when they interact with an inhibitor. In a genetic network, genes are active when they are expressed (experimentally detected as over-expression) and inactive when they are repressed (experimentally detected as under-expression). In the protein-protein interaction (PPI) networks, some proteins carry out their interactions only when they are phosphorylated, thus becoming active, and vice versa by dephosphorylation. By default, neutral proteins remain unaffected, neither active nor inactive, for a particular phenotype.

#### 2. TPMS modelling of phenotypes.

The Therapeutic Performance Mapping System (TPMS) is a tool that creates mathematical models of a drug/pathology response to explain a clinical outcome or phenotype[1,3–8]. These models find MoAs that explain how a *Stimulus* (i.e. proteins activated or inactivated by a drug) produces a *Response* (i.e. proteins activated or inactivated in a phenotype). As example of use, here we apply TPMS to the drug-indication pair sacubitril/valsartan and HF: for the drug we retrieve the sacubitril/valsartan targets from DrugBank [9], PubChem[10], STITCH[11], SuperTarget[12] and hand curated literature revision. Afterward, we consider the proteins whose modulations have been associated with HF from the BED[1,2]. Finally, after applying the TPMS function, we obtain a set of connected proteins (subnetworks) with associated activities, each subnetwork with a potential explanation of the molecular mechanism of the drug in agreement with what has been previously described (i.e. a potential MoA).

##### 2.1. Building the Human protein network (HPN)

To apply the TPMS function and create mathematical models of MoAs, we first need to develop the HPN. In this study, we use a PPI network created from the integration of public and private databases: KEGG[13], BioGRID[14], IntAct[15], REACTOME[16], TRRUST[17], and HPRD[18]. In addition, we include information extracted from scientific literature, which is manually curated and used to trim the network.

##### 2.2. Defining restrictions from gene expression data

In order to train and validate the models, it is necessary to obtain a collection of restrictions that are defined as the "true set". The basis restrictions are obtained from HPRD[18], DIP[19], TRRUST[17], INTACT[15], REACTOME[16], BIOGRID[14], SIDER[20] and DrugBank[9]. They help to indicate what proteins are active or inactive specifically for a human particular phenotype. Additionally, we include specific restrictions derived from gene expression data as defined by the user (i.e. adding specific information in our test example from changes of expression induced by sacubitril/valsartan, or transcriptomic data on HF phenotypes). Hence, we have used the GSE57345 dataset[21] as in *Iborra-Egea et al.*[3] We calculate the fold change of genes associated with the HPN and map the gene expression data as activated or inhibited proteins (active if they are produced by over-expressed genes, and inactive -inhibited- if produced by under-expressed genes).

##### 2.3. Description of the mathematical models

The algorithm of TPMS to generate the models is similar to a Multilayer Perceptron of an Artificial Neural Network over the HPN (where neurons are the proteins and the edges of the network are used to transfer the information). We consider as input signals the values of activation (+1) and inactivation (-1) of the targets of a drug. The output results are then the values of activation and inactivation of the proteins defining the phenotype (as retrieved from the BED), named effectors. We limit the network by considering only interactions that connect drug targets with protein effectors in a maximum of three steps. The parameters to solve are the weights associated to the links between two nodes ($\omega_l$). Each node of the protein network receives as input the output of the connected nodes in the direction flow from targets to effectors, weighted by each link weight ($\omega_l$). The sum of inputs is transformed by a hyperbolic tangent function to generate the score of the node (neuron), which become the "output signal" of the current node towards the nodes. Details of the approach are shown in **Figure 1a**, where $n_5$ is linked to $n_1$ and $n_2$. The output signal of $n_5$ is $n_5 = \tanh(n_1 \cdot \omega_{1-5} + n_2 \cdot \omega_{2-5})$. We obtain the $\omega_l$ parameters by optimization, using a Stochastic Optimization Method based on Simulated Annealing[22], such that the values of the nodes in the effectors are the closest to their expected value. The models are trained by using the restrictions defined by the BED and the specific data set by the user (i.e. the GSE57345 dataset[21] of gene-expression as in *Iborra-Egea et al.*[3] mentioned above). The iterative process of optimization usually requires between $10^6$ and $10^9$ iterations, until satisfying at least the 80% of the restrictions and the values of the effectors. However, the number of $\omega_l$ parameters is very high (between 100,000 and 400,000 depending on the size of the subnetwork) and the size of the collection of restrictions (approximately $10^7$) is usually not enough to find a unique solution. Consequently, the TPMS approach finds a set of potential solutions. We rank all solutions by the number of restrictions satisfied and select the top 200 solutions satisfying the largest number, including the expected values of the effectors. These solutions represent 200 potential MoAs of the drug, which we assume equally acceptable and with the same probability of occurrence. Here, we hypothesize that these solutions represent different cells, while combinations of them would correspond to different patients. Hence, 200 prototype or representative mathematical solutions can be considered for an individual and personalized approach (see **Figure 1b**).

#### 3. Measures to compare sets of MoAs

TPMS returns a set of MoAs describing potential relationships between the targets of a drug and the biological effectors of a disease. We hypothesize that TPMS solutions represent different MoAs in cells and consequently as combinations in a population of patients. Therefore, to understand the relationships between all potential mechanisms we need to define measures of comparison between different sets of solutions. Here, we define several measures in order to study and compare sets of MoAs from different views.

##### 3.1. Intensity of the response

We defined the “intensity” of the response as a measure to qualify a MoA and compare it with others. The intensity is defined as a pair: 1) the number of protein effectors (#) achieving an expected signal sign; and 2) a measure of the strength of the output signal of the effectors (i.e. a global measure of the output signal, named TSignal). Assuming $y_i$ as the value achieved by a protein effector “i”, while $v_i$ is the effector sign according to the BED (active or inactive) and $n$ is the total number of effectors described for a phenotype, we define:

- **Number of effectors achieving the expected sign**: We expect that a drug will revert the conditions of a disease phenotype, while it may reach the effectors of an adverse event. Consequently, a drug should inactivate the active protein effectors of a pathology-phenotype and activate the inactive ones, but it could activate/inhibit other adverse event effectors with the same sign as described in the BED. Using Dirac’s $\delta$ (i.e. $\delta(0)=1$, and zero otherwise), for drug indications the formula is:

  $$\#_{indication} = \sum_{i=1}^{n} \delta\left(v_i + \frac{y_i}{|y_i|}\right) \qquad \textbf{[Equation 1a]}$$

  So, in the case of the disease effectors we only count the effectors with a BED value of opposite sign to the signal arriving from the drug.

  However, for adverse events the formula changes because we count the adverse event effectors that are affected by the drug, and therefore the signal arriving from the drug has the same sign as the BED value of the effector:

  $$\#_{adverse\ event} = \sum_{i=1}^{n} \delta\left(v_i - \frac{y_i}{|y_i|}\right) \qquad \textbf{[Equation 1b]}$$

- **TSignal**: The average of the output signals of the protein effectors with the correct sign considered as positive signal, and the ones with the incorrect sign considered as negative signal. For a drug affecting the phenotype of a disease, this implies that $v_i$ and $y_i$ have opposite sign and we need to change the sign:

  $$TSignal_{indication} = -\frac{1}{n}\sum_{i=1}^{n} v_i y_i \qquad \textbf{[Equation 2a]}$$

  On the contrary, for testing if a drug introduces adverse events we check if the output signal has the same sign as the effector of the disease phenotype, and therefore TSignal is defined as:

  $$TSignal_{adverse\ event} = \frac{1}{n}\sum_{i=1}^{n} v_i y_i \qquad \textbf{[Equation 2b]}$$

##### 3.2. Distance between two sets of MoAs

We define the *distance* between two or more sets of MoAs in order to determine their similarity. To compute the distance, we use a modified Hausdorff distance (MHD) introduced by Dubuisson and Jain.[23] We use the distance measures between two (finite) point sets A and B:

$$\text{For } a \in A, \qquad d(a,B) \coloneqq \min_{b\in B} d(a,b),$$

$$\text{and} \qquad d_A(B) \coloneqq \frac{1}{|A|}\sum_{a\in A} d(a,B),$$

Where |A| is the number of elements in A, d(·,·) is the Euclidean distance and “a” and “b” are n-tuples of the activities (output signals) of the nodes of two MoAs (a in A and b in B). Then, we define the MHD as:

$$d_{\mathrm{MHD}}(A,B) \coloneqq \max\left( d_A(B), d_B(A) \right) \qquad \textbf{[Equation 3]}$$

Note that the MHD is a semimetric and not a metric, since the triangular inequality does not hold.

##### 3.3. Potential biomarkers extracted from MoAs

To identify potential biomarkers and stratify patients according to the drug response and adverse effects, we group the MoAs. For HF, MoAs are ranked by their TSignal and split in four quartiles: the first quartile (top 25%) contains MoAs with higher intensity of the response (TSignal), which in turn reduces the values of the effectors associated with a disease phenotype (we name them as “Low”-disease MoAs). On the contrary, the fourth quartile (bottom 25%) collects MoAs with lower intensity of response (thus, we named as “High”-disease MoAs). On the other hand, for MD, the first quartile (top 25%) contains MoAs with higher intensity, which in this case, as an adverse event, it increases the values of the effectors associated to the comorbidity (we name them as High-adverseEvent MoAs). The fourth quartile (bottom 25%) collects MoAs with lower intensity of response (thus, we named as Low- adverseEvent MoAs).

We use the comparison between both groups of High- and Low- MoAs to identify the *best-classifier proteins*, specific proteins helping us to infer biological associations and distinguish the responses of drugs on a population (i.e. potential biomarkers). Best-classifier proteins (single or pairs) are the proteins from the MoAs that can divide better the samples between High- and Low-groups. These classifiers are determined by a Data-Science strategy based on the use of a set of Feature Selection algorithms combined with Base Classifiers. The feature selection used for single proteins was brute force[24], so analysing one feature or protein at a time, while for protein pairs the following Base Classifiers were used: elastic net (Zou, H. and T. Hastie. Regularization and variable selection via the elastic net. Journal of the Royal Statistical Society, Series B, Vol. 67, No. 2, pp. 301–320, 2005.); entropy and correlation (Pedregosa et al., JMLR 12, pp. 2825-2830, 2011); LASSO (Tibshirani, Robert. 1996. "Regression Shrinkage and Selection via the lasso". Journal of the Royal Statistical Society. Series B (methodological) 58 (1). Wiley: 267–88. http://www.jstor.org/stable/2346178); random forest (Ho, Tin Kam (1995). Random Decision Forests (PDF). Proceedings of the 3rd International Conference on Document Analysis and Recognition, Montreal, QC, 14–16 August 1995. pp. 278–282.); GLM random sets (Madsen, Henrik, and Thyregod, Poul (2011). Introduction to General and Generalized Linear Models. Chapman & Hall/CRC. ISBN 978-1-4200-9155-7.); ReliefF (Kira and Rendell in 1992 Kira, Kenji and Rendell, Larry (1992).); Ridge regression (Feature Selection based on the Bhattacharyya Distance Guorong Xuan et al. "Feature Selection based on the Bhattacharyya Distance.".January 2006); simple regression (Keinosuke Fukunaga, Introduction to statistical pattern recognition (2nd ed.) Academic Press Professional, Inc. San Diego, CA, USA 1990 ISBN:0-12-269851-7.); Wilcoxon test (A Critical Assessment of Feature Selection Methods for Biomarker Discovery in Clinical Proteomics. Christin C1, Hoefsloot HC, Smilde AK, Hoekman B, Suits F, Bischoff R, Horvatovich P.); Wilcoxon test with correlation (A Critical Assessment of Feature Selection Methods for Biomarker Discovery in Clinical Proteomics. Christin C1, Hoefsloot HC, Smilde AK, Hoekman B, Suits F, Bischoff R, Horvatovich P.).Several base classifiers were applied to distinguish the two groups using the selected features: optimal threshold; linear regression[25]; Multilayer Perceptron Network[26]; Generalized Linear Model[27]; elastic net[28]; optimal quadratic threshold[29]. Best-classifier proteins will be the 200 proteins (or pair of proteins) that after k-fold cross-validation (k=10)[30] have the best balanced accuracy[31] of the classification. Best-classifier proteins are strongly related to the intensity of a response and are differently distributed between Low- and High-group MoAs. Assuming the hypothesis that the selected MoAs are representative of individual prototype patients, these proteins can be used as biomarkers to classify a cohort of patients by the activity or absence of activity of the protein.

Each best-classifier protein has a distribution of signal values corresponding to the Low-group MoAs and another corresponding to the High-group MoAs. We use a Mann-Whitney *U* test to compare the two distributions of signal values and select those proteins that have a significantly different distribution (p-value< 0.01), having an average output signal in Low-HF with opposite sign to the average output signal in High-HF (i.e. positive vs. negative or vice versa). We name these as *differential best-classifier proteins*. By following this strategy, we can identify two groups of differential best-classifier proteins: those active in Low-group and inactive in High-group, and those active in High-grout but inactive in Low-group.

### Figures

(a) (b)

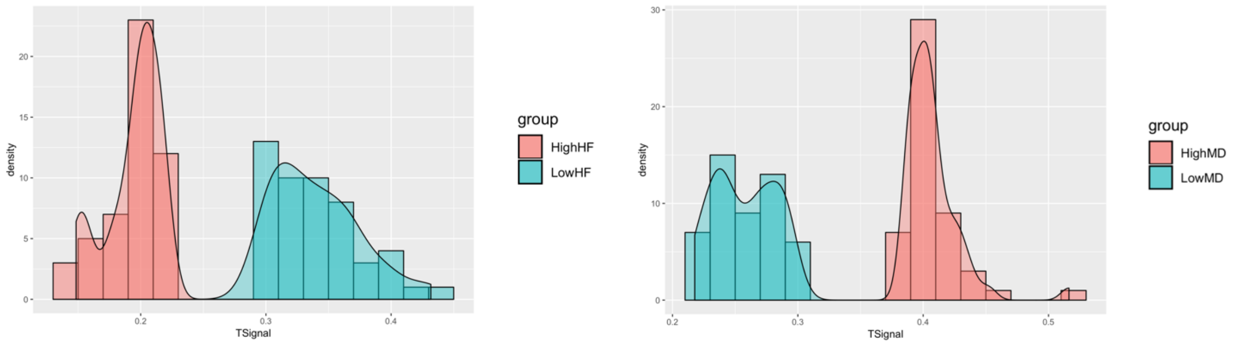


**Supplementary Figure 1:** Histogram of the number of models belonging to High- and Low- (HF in **(a)** and MD in **(b)**) in a range of TSignal values.

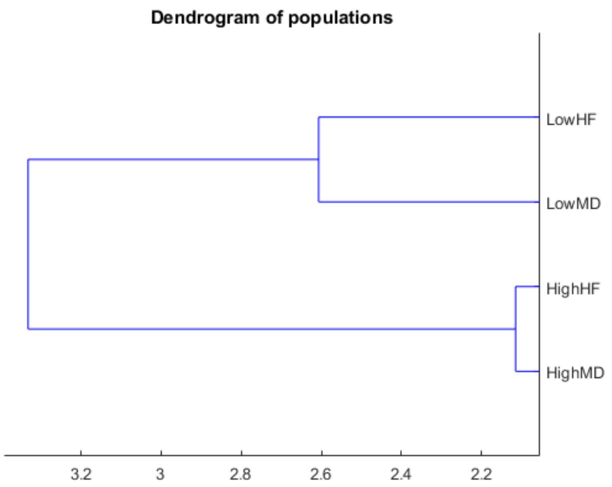


**Supplementary Figure 2**: Dendrogram plot of the pairwise modified Hausdorff distance (MHD) between the four groups of mechanisms of action (MoAs): LowHF, HighHF, LowMD, HighMD.

(a) (b)

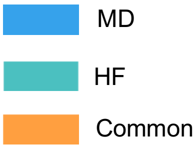

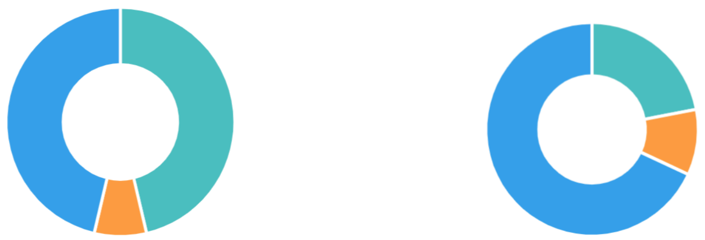

| N common | N1 | N2 | N total | Odds Ratio | P value |
|---|---|---|---|---|---|
| 36 | 262 | 262 | 13090 | 8.882 | 3.1E-20 |

| N common | N1 | N2 | N total | Odds Ratio | P value |
|---|---|---|---|---|---|
| 19 | 61 | 149 | 8520 | 28.984 | 2.0E-19 |

**Supplementary Figure 3**: Screenshots of the results of the genetic and functional overlap between the GUILDify subnetworks of Heart Failure (HF) and Macular Degeneration (MD). The **(a)** and **(b)** sections show the results of the genetic and functional overlap respectively.

### Tables

**Supplementary Table 1**: Differential best-classifier proteins with opposite signal in Low-HF (LHF) and High-HF (HHF). "+" stands for active, while "-" stands for inactive. Highlighted cells correspond to proteins that are part of the Top-HF ∪ Top-MD ∪ Top-Drug set, the top-scoring proteins according to GUILDify

| | Uniprot ID | Gene symbol | Gene name | $\langle LHF \rangle$ | $\langle HHF \rangle$ | $\sqrt{\lvert LMDxHMD \rvert}$ | Adjusted P-value |
|---|---|---|---|---|---|---|---|
| **LHF+ HHF-** | Q96F07 | CYFIP2 | Cytoplasmic FMR1-interacting protein 2 | 0.110 | -0.278 | 0.175 | 6.388E-07 |
| | P55160 | NCKAP1L | Nck-associated protein 1-like {ECO:0000305} | 0.110 | -0.278 | 0.175 | 6.388E-07 |
| | Q7L576 | CYFIP1 | Cytoplasmic FMR1-interacting protein 1 | 0.110 | -0.278 | 0.175 | 6.388E-07 |
| | Q9NYB9 | ABI2 | Abl interactor 2 | 0.110 | -0.278 | 0.175 | 6.388E-07 |
| | Q9Y2A7 | NCKAP1 | Nck-associated protein 1 | 0.110 | -0.278 | 0.175 | 6.388E-07 |
| | P50052 | AGTR2 | Type-2 angiotensin II receptor | 0.205 | -0.013 | 0.051 | 1.852E-05 |
| **LHF- HHF+** | **P28482** | **MAPK1** | **Mitogen-activated protein kinase 1** | -0.710 | 0.479 | 0.584 | 1.854E-14 |
| | **P27361** | **MAPK3** | **Mitogen-activated protein kinase 3** | -0.313 | 0.962 | 0.549 | 6.366E-15 |
| | P47900 | P2RY1 | P2Y purinoceptor 1 | -0.322 | 0.605 | 0.441 | 9.855E-10 |
| | Q92558 | WASF1 | Wiskott-Aldrich syndrome protein family member 1 | -0.580 | 0.309 | 0.424 | 8.494E-13 |
| | Q9Y6W5 | WASF2 | Wiskott-Aldrich syndrome protein family member 2 | -0.580 | 0.309 | 0.424 | 8.494E-13 |
| | O00401 | WASL | Neural Wiskott-Aldrich syndrome protein | -0.580 | 0.309 | 0.424 | 8.494E-13 |
| | **P02751** | **FN1** | **Fibronectin** | -0.476 | 0.224 | 0.327 | 2.152E-09 |
| | O60229 | KALRN | Kalirin {ECO:0000250\|UniProtKB:P97924} | -0.529 | 0.185 | 0.313 | 3.239E-05 |
| | Q8TEW0 | PARD3 | Partitioning defective 3 homolog | -0.279 | 0.255 | 0.267 | 3.577E-09 |
| | P41743 | PRKCI | Protein kinase C iota type | -0.279 | 0.255 | 0.267 | 3.577E-09 |
| | Q13576 | IQGAP2 | Ras GTPase-activating-like protein IQGAP2 | -0.279 | 0.255 | 0.267 | 3.577E-09 |
| | O75914 | PAK3 | Serine/threonine-protein kinase PAK 3 | -0.279 | 0.255 | 0.267 | 3.577E-09 |
| | Q9P286 | PAK5 | Serine/threonine-protein kinase PAK 5 {ECO:0000305} | -0.279 | 0.255 | 0.267 | 3.577E-09 |
| | O96013 | PAK4 | Serine/threonine-protein kinase PAK 4 | -0.279 | 0.255 | 0.267 | 3.577E- |

| | | | | | | | |
|---|---|---|---|---|---|---|---|
| | | | | | | | 09 |
| | Q86VI3 | IQGAP3 | Ras GTPase-activating-like protein IQGAP3 | -0.279 | 0.255 | 0.267 | 3.577E-09 |
| | Q9NPB6 | PARD6A | Partitioning defective 6 homolog alpha | -0.279 | 0.255 | 0.267 | 3.577E-09 |
| | Q16584 | MAP3K11 | Mitogen-activated protein kinase kinase kinase 11 | -0.279 | 0.255 | 0.267 | 3.577E-09 |
| | Q9NQU5 | BUB1B-PAK6; PAK6 | Serine/threonine-protein kinase PAK 6 | -0.279 | 0.255 | 0.267 | 3.577E-09 |
| | Q9BYG5 | PARD6B | Partitioning defective 6 homolog beta | -0.279 | 0.255 | 0.267 | 3.577E-09 |
| | Q9BYG4 | PARD6G | Partitioning defective 6 homolog gamma | -0.279 | 0.255 | 0.267 | 3.577E-09 |
| | P84095 | RHOG | Rho-related GTP-binding protein RhoG | -0.449 | 0.156 | 0.265 | 1.935E-09 |
| | Q96PN6 | ADCY10 | Adenylate cyclase type 10 | -0.488 | 0.144 | 0.265 | 2.421E-09 |
| | Q96JJ3 | ELMO2 | Engulfment and cell motility protein 2 | -0.396 | 0.137 | 0.233 | 1.935E-09 |
| | Q15759 | MAPK11 | Mitogen-activated protein kinase 11 | -0.883 | 0.056 | 0.223 | 6.366E-15 |
| | O15264 | MAPK13 | Mitogen-activated protein kinase 13 | -0.884 | 0.045 | 0.198 | 6.366E-15 |
| | P53778 | MAPK12 | Mitogen-activated protein kinase 12 | -0.884 | 0.045 | 0.198 | 6.366E-15 |
| | P54764 | EPHA4 | Ephrin type-A receptor 4 | -0.139 | 0.233 | 0.180 | 6.388E-07 |
| | Q99755 | PIP5K1A | Phosphatidylinositol 4-phosphate 5-kinase type-1 alpha | -0.110 | 0.278 | 0.175 | 6.388E-07 |
| | Q9UQB8 | BAIAP2 | Brain-specific angiogenesis inhibitor 1-associated protein 2 | -0.110 | 0.278 | 0.175 | 6.388E-07 |
| | O14986 | PIP5K1B | Phosphatidylinositol 4-phosphate 5-kinase type-1 beta | -0.110 | 0.278 | 0.175 | 6.388E-07 |
| | Q9Y5S8 | NOX1 | NADPH oxidase 1 | -0.110 | 0.278 | 0.175 | 6.388E-07 |
| | P46734 | MAP2K3 | Dual specificity mitogen-activated protein kinase kinase 3 | -0.110 | 0.278 | 0.175 | 6.388E-07 |
| | Q15080 | NCF4 | Neutrophil cytosol factor 4 | -0.110 | 0.278 | 0.175 | 6.388E-07 |
| | Q9HBY0 | NOX3 | NADPH oxidase 3 | -0.110 | 0.278 | 0.175 | 6.388E-07 |
| | Q9UPY6 | WASF3 | Wiskott-Aldrich syndrome protein family member 3 | -0.110 | 0.278 | 0.175 | 6.388E-07 |
| | P52564 | MAP2K6 | Dual specificity mitogen-activated protein kinase kinase 6 | -0.110 | 0.278 | 0.175 | 6.388E-07 |
| | O14733 | MAP2K7 | Dual specificity mitogen-activated protein kinase kinase 7 | -0.110 | 0.278 | 0.175 | 6.388E-07 |
| | Q9Y4K3 | TRAF6 | TNF receptor-associated factor 6 | -0.097 | 0.285 | 0.166 | 1.917E-06 |
| | P19878 | NCF2 | Neutrophil cytosol factor 2 | -0.042 | 0.278 | 0.108 | 3.050E-05 |

**Supplementary Table 2**: Top 10 gene Ontology functions enriched from best-classifier proteins with opposite signal in Heart Failure (HF) MoAs. Functional enrichment analysis from FuncAssociate.

| | **Low-HF active / High-HF inactive** | | | **Low-HF inactive / High-HF active** | | | **Overlapped functions** | | |
|---|---|---|---|---|---|---|---|---|---|
| | **GO name** | **LOD** | **P-val.** | **GO name** | **LOD** | **P-val.** | **GO name** | **LOD** | **P-val.** |
| **1** | SCAR complex | 3.89 | <0.00050 | 1-phosphatidylinositol-3-phosphate 5-kinase activity | 3.41 | 0.01700 | Rac protein signal transduction | 2.54 | <0.00050 |
| **2** | positive regulation of Arp2/3 complex-mediated actin nucleation | 3.64 | <0.00050 | 1-phosphatidylinositol-5-kinase activity | 3.41 | 0.01700 | vascular endothelial growth factor receptor signaling pathway | 2.31 | <0.00050 |

| | | | | | | | | | |
|---|---|---|---|---|---|---|---|---|---|
| **3** | positive regulation of neurotrophin TRK receptor signaling pathway | 3.49 | 0.00150 | phosphatidylinositol-3,4-bisphosphate 5-kinase activity | 2.94 | 0.04000 | immune response-regulating cell surface receptor signaling pathway involved in phagocytosis | 1.95 | <0.00050 |
| **4** | regulation of neurotrophin TRK receptor signaling pathway | 3.36 | <0.00050 | proteolysis in other organism | 2.73 | 0.00250 | Fc-gamma receptor signaling pathway involved in phagocytosis | 1.95 | <0.00050 |
| **5** | positive regulation of actin nucleation | 3.27 | <0.00050 | MAP kinase kinase activity | 2.60 | <0.00050 | Fc receptor mediated stimulatory signaling pathway | 1.95 | <0.00050 |
| **6** | regulation of Arp2/3 complex-mediated actin nucleation | 3.23 | <0.00050 | NADPH oxidase complex | 2.58 | <0.00050 | Fc-gamma receptor signaling pathway | 1.94 | <0.00050 |
| **7** | dendrite extension | 3.16 | 0.00350 | DNA damage induced protein phosphorylation | 2.53 | 0.00450 | Fc receptor signaling pathway | 1.74 | <0.00050 |
| **8** | regulation of actin nucleation | 2.96 | <0.00050 | MAP kinase activity | 2.52 | <0.00050 | Ras protein signal transduction | 1.79 | <0.00050 |
| **9** | filopodium tip | 2.84 | 0.01200 | superoxide-generating NADPH oxidase activity | 2.34 | 0.00600 | regulation of actin cytoskeleton organization | 1.62 | 0.00072 |
| **10** | developmental cell growth | 2.42 | 0.00150 | superoxide anion generation | 2.25 | 0.00850 | lamellipodium | 1.71 | 0.00086 |

**Supplementary Table 3**: Differential best-classifier proteins with opposite signal in Low-MD (LMD) and High-MD (HMD). "+" stands for active, while "-" stands for inactive. Highlighted cells correspond to proteins that are part of the Top-HF ∪ Top-MD ∪ Top-Drug set, the top-scoring proteins according to GUILDify

| | **Uniprot ID** | **Gene symbol** | **Gene name** | $\langle LMD \rangle$ | $\langle HMD \rangle$ | $\sqrt{\lvert LMDxHMD \rvert}$ | **Adjusted P-value** |
|---|---|---|---|---|---|---|---|
| **LMD+ HMD-** | Q9Y4H2 | IRS2 | Insulin receptor substrate 2 | 0.583 | -0.414 | 0.491 | 1.297E-13 |
| | O43639 | NCK2 | Cytoplasmic protein NCK2 | 0.623 | -0.355 | 0.471 | 5.744E-11 |
| | Q13153 | PAK1 | Serine/threonine-protein kinase PAK 1 {ECO:0000303\|PubMed:8805275} | 0.233 | -0.817 | 0.437 | 2.266E-12 |
| | P30530 | AXL | Tyrosine-protein kinase receptor UFO | 0.476 | -0.362 | 0.415 | 5.509E-16 |
| | P42081 | CD86 | T-lymphocyte activation antigen CD86 | 0.428 | -0.356 | 0.391 | 2.073E-08 |
| | P18825 | ADRA2C | Alpha-2C adrenergic receptor | 0.226 | -0.568 | 0.358 | 2.079E-10 |
| | Q13177 | PAK2 | Serine/threonine-protein kinase PAK 2 | 0.249 | -0.439 | 0.330 | 3.753E-09 |
| | P54762 | EPHB1 | Ephrin type-B receptor 1 | 0.144 | -0.685 | 0.314 | 2.916E-14 |
| | P15498 | VAV1 | Proto-oncogene vav | 0.392 | -0.192 | 0.274 | 8.020E-05 |
| | P06241 | FYN | Tyrosine-protein kinase Fyn | 0.589 | -0.127 | 0.274 | 7.322E-15 |
| | **O75787** | **ATP6AP2** | **V-ATPase M8.9 subunit** | 0.407 | -0.160 | 0.255 | 2.741E-08 |
| | **P01583** | **IL1A** | **Interleukin-1 alpha** | 0.125 | -0.396 | 0.222 | 2.087E-12 |
| | **P06748** | **NPM1** | **Nucleophosmin** | 0.374 | -0.116 | 0.208 | 2.266E-12 |

| | | | | | | | |
|---|---|---|---|---|---|---|---|
| | **Q02297** | **NRG1** | **Pro-neuregulin-1, membrane-bound isoform** | 0.670 | -0.064 | 0.207 | 5.208E-14 |
| | P15941 | MUC1 | Mucin-1 subunit alpha | 0.085 | -0.479 | 0.202 | 1.676E-11 |
| | **P18084** | **ITGB5** | **Integrin beta-5** | 0.498 | -0.079 | 0.199 | 1.214E-15 |
| | P03372 | ESR1 | Estrogen receptor | 0.096 | -0.294 | 0.169 | 6.103E-08 |
| | P01138 | NGF | Beta-nerve growth factor | 0.211 | -0.124 | 0.162 | 6.954E-07 |
| | P43405 | SYK | Tyrosine-protein kinase SYK | 0.075 | -0.310 | 0.152 | 1.618E-07 |
| | Q08722 | CD47 | Leukocyte surface antigen CD47 | 0.082 | -0.277 | 0.151 | 8.239E-07 |
| | P54764 | EPHA4 | Ephrin type-A receptor 4 | 0.336 | -0.065 | 0.148 | 4.859E-08 |
| | Q9BYF1 | ACE2 | Processed angiotensin-converting enzyme 2 | 0.565 | -0.039 | 0.148 | 7.333E-15 |
| | P10275 | AR | Androgen receptor | 0.438 | -0.045 | 0.141 | 1.014E-11 |
| | P38398 | BRCA1 | Breast cancer type 1 susceptibility protein | 0.043 | -0.365 | 0.125 | 9.363E-08 |
| | P35354 | PTGS2 | Prostaglandin G/H synthase 2 | 0.034 | -0.396 | 0.116 | 2.482E-12 |
| | Q9Y478 | PRKAB1 | 5'-AMP-activated protein kinase subunit beta-1 | 0.374 | -0.034 | 0.113 | 5.744E-11 |
| | P14770 | GP9 | Platelet glycoprotein IX | 0.034 | -0.306 | 0.102 | 1.190E-08 |
| | P14138 | EDN3 | Endothelin-3 | 0.023 | -0.239 | 0.074 | 3.509E-06 |
| **LMD-HMD+** | **P02675** | **FGB** | **Fibrinogen beta chain** | -0.778 | 0.654 | 0.713 | 3.040E-14 |
| | O60674 | JAK2 | Tyrosine-protein kinase JAK2 | -0.811 | 0.279 | 0.476 | 2.749E-16 |
| | **P04085** | **PDGFA** | **Platelet-derived growth factor subunit A** | -0.359 | 0.622 | 0.473 | 1.263E-07 |
| | Q05586 | GRIN1 | Glutamate receptor ionotropic, NMDA 1 | -0.381 | 0.565 | 0.464 | 1.049E-15 |
| | **P05230** | **FGF1** | **Fibroblast growth factor 1** | -0.219 | 0.734 | 0.401 | 1.528E-14 |
| | Q15768 | EFNB3 | Ephrin-B3 | -0.149 | 0.835 | 0.353 | 8.307E-13 |
| | Q14451 | GRB7 | Growth factor receptor-bound protein 7 | -0.181 | 0.679 | 0.351 | 5.106E-13 |
| | P08581 | MET | Hepatocyte growth factor receptor | -0.124 | 0.828 | 0.321 | 2.615E-13 |
| | Q08289 | CACNB2 | Voltage-dependent L-type calcium channel subunit beta-2 | -0.351 | 0.238 | 0.289 | 5.549E-09 |
| | P63244 | RACK1 | Receptor of activated protein C kinase 1, N-terminally processed | -0.395 | 0.206 | 0.285 | 4.410E-08 |
| | Q00987 | MDM2 | E3 ubiquitin-protein ligase Mdm2 | -0.458 | 0.166 | 0.275 | 2.789E-08 |
| | P32004 | L1CAM | Neural cell adhesion molecule L1 | -0.466 | 0.118 | 0.235 | 2.519E-12 |
| | P15391 | CD19 | B-lymphocyte antigen CD19 | -0.272 | 0.171 | 0.216 | 4.123E-08 |
| | P07948 | LYN | Tyrosine-protein kinase Lyn | -0.109 | 0.408 | 0.211 | 3.099E-04 |
| | O14745 | SLC9A3R1 | Na(+)/H(+) exchange regulatory cofactor NHE-RF1 | -0.172 | 0.224 | 0.196 | 4.627E-07 |
| | O43559 | FRS3 | Fibroblast growth factor receptor substrate 3 | -0.091 | 0.317 | 0.170 | 3.717E-08 |
| | P43146 | DCC | Netrin receptor DCC | -0.392 | 0.070 | 0.165 | 5.835E-04 |
| | P62158 | CALM1 ; CALM2 ; CALM3 | Calmodulin-1 {ECO:0000312\|HGNC:HGNC:1442} | -0.455 | 0.054 | 0.156 | 1.670E-10 |
| | **P42574** | **CASP3** | **Caspase-3 subunit p12** | -0.034 | 0.656 | 0.149 | 8.050E-08 |
| | P42684 | ABL2 | Abelson tyrosine-protein kinase 2 | -0.362 | 0.045 | 0.128 | 1.676E-11 |
| | P17081 | RHOQ | Rho-related GTP-binding protein RhoQ | -0.362 | 0.045 | 0.128 | 1.676E-11 |
| | Q13905 | RAPGEF1 | Rap guanine nucleotide exchange factor 1 | -0.187 | 0.080 | 0.122 | 2.094E-04 |
| | **P05155** | **SERPING1** | **Plasma protease C1 inhibitor** | -0.023 | 0.362 | 0.091 | 1.014E-11 |
| | Q92793 | CREBBP | CREB-binding protein | -0.506 | 0.015 | 0.089 | 4.511E- |

| | | | | | | | |
|---|---|---|---|---|---|---|---|
| | | | | | | | 11 |
| | P07585 | DCN | Decorin | -0.023 | 0.351 | 0.089 | 2.430E-11 |
| | P12830 | CDH1 | Cadherin-1 | -0.503 | 0.011 | 0.076 | 1.487E-14 |
| | Q07157 | TJP1 | Tight junction protein ZO-1 | -0.407 | 0.011 | 0.068 | 2.640E-12 |
| | Q92990 | GLMN | Glomulin | -0.294 | 0.011 | 0.058 | 2.056E-07 |
| | P55075 | FGF8 | Fibroblast growth factor 8 | -0.011 | 0.238 | 0.052 | 1.884E-05 |

**Supplementary Table 4**: Top 10 gene Ontology functions enriched from best-classifier proteins with opposite signal in Macular Degeneration (MD) MoAs. Functional enrichment analysis from FuncAssociate.

| | **Low-MD active / High-MD inactive** | | | **Low-MD inactive / High-MD active** | | | **Overlapped functions** | | |
|---|---|---|---|---|---|---|---|---|---|
| | **GO name** | **LOD** | **P-val.** | **GO name** | **LOD** | **P-val.** | **GO name** | **LOD** | **P-val.** |
| **1** | dendritic spine development | 2.41 | 0.00150 | dorsal/ventral axon guidance | 3.07 | 0.01950 | phosphatidylinositol-4,5-bisphosphate 3-kinase activity | 1.89 | <0.00050 |
| **2** | positive regulation of vascular endothelial growth factor production | 2.04 | 0.03000 | fibroblast growth factor receptor binding | 2.06 | 0.02000 | phosphatidylinositol bisphosphate kinase activity | 1.87 | <0.00050 |
| **3** | regulation of intracellular estrogen receptor signaling pathway | 2.00 | 0.00150 | platelet-derived growth factor receptor signaling pathway | 1.95 | 0.03350 | phosphatidylinositol 3-kinase activity | 1.84 | <0.00050 |
| **4** | regulation of systemic arterial blood pressure | 1.97 | 0.03300 | non-membrane spanning protein tyrosine kinase activity | 1.88 | 0.04000 | phosphatidylinositol phosphorylation | 1.72 | <0.00050 |
| **5** | regulation of vascular endothelial growth factor production | 1.96 | 0.04450 | growth factor receptor binding | 1.85 | <0.00050 | single-organism cellular process | 1.53 | <0.00050 |
| **6** | peptide hormone processing | 1.96 | 0.04450 | regulation of blood coagulation | 1.68 | 0.01050 | lipid phosphorylation | 1.67 | <0.00050 |
| **7** | phosphotyrosine binding | 1.91 | 0.04900 | regulation of hemostasis | 1.68 | 0.01050 | positive regulation of protein kinase B signaling | 1.60 | <0.00050 |
| **8** | neutrophil chemotaxis | 1.90 | 0.05000 | regulation of coagulation | 1.66 | 0.01050 | biological regulation | 1.42 | <0.00050 |
| **9** | regulation of vasoconstriction | 1.89 | 0.00150 | regulation of phosphatidylinositol 3-kinase signaling | 1.58 | 0.02350 | protein binding | 1.41 | <0.00050 |
| **10** | vascular endothelial growth factor receptor signaling pathway | 1.83 | <0.00050 | response to toxic substance | 1.53 | 0.00350 | regulation of response to stimulus | 1.24 | <0.00050 |

**Supplementary Table 5**: Modified Hausdorff distance between the 4 groups of MoAs defined.

| | LowMD | HighMD | HighHF | LowHF |
|---|---|---|---|---|
| LowMD | 0 | 4.00226983 | 2.7537393 | 2.6068664 |
| HighMD | 4.00226983 | 0 | 2.1150102 | 2.55445687 |
| HighHF | 2.7537393 | 2.1150102 | 0 | 4.01919608 |

| | | | | |
|---|---|---|---|---|
| LowHF | 2.6068664 | 2.55445687 | 4.01919608 | 0 |

**Supplementary Table 6**: Mean Euclidean distance between each one of the points of every group of MoAs and its centre.

| | Mean distance from center |
|---|---|
| LowMD | 3.137818031 |
| HighMD | 3.171767895 |
| HighHF | 3.298746704 |
| LowHF | 3.523965485 |

**Supplementary Table 7**: Number of common MoAs between the 4 groups of MoAs defined.

| | LowMD | HighMD | HighHF | LowHF |
|---|---|---|---|---|
| LowMD | 50 | 0 | 9 | 13 |
| HighMD | 0 | 50 | 17 | 12 |
| HighHF | 9 | 17 | 50 | 0 |
| LowHF | 13 | 12 | 0 | 50 |

**Supplementary Table 8**: Intersection of several set of proteins defined with GUILDify with the best-classifier proteins (BCP) obtained from the TPMS analysis. The p-values are calculated using a Fisher's exact test. The p-values above 0.05 are remarked in red.

| **Sets of proteins** | **# LHF+ HHF-** | **P-value** | **# LHF- HHF+** | **P-value** | **# LMD+ HMD-** | **P-value** | **# LMD- HMD+** | **P-value** |
|---|---|---|---|---|---|---|---|---|
| Drug seeds | 0 | 1.00E+00 | 0 | 1.00E+00 | 0 | 1.00E+00 | 0 | 1.00E+00 |
| Top-Drug | 0 | 1.00E+00 | 0 | 1.00E+00 | 2 | 1.11E-01 | 0 | 1.00E+00 |
| HF seeds | 0 | 1.00E+00 | 3 | 6.32E-03 | 1 | 2.32E-01 | 1 | 2.39E-01 |
| Top-HF | 0 | 1.00E+00 | 3 | 4.34E-02 | 2 | 1.02E-01 | 1 | 4.35E-01 |
| MD seeds | 0 | 1.00E+00 | 1 | 4.02E-01 | 2 | 4.81E-02 | 5 | 2.76E-05 |
| Top-MD | 0 | 1.00E+00 | 1 | 5.60E-01 | 3 | 1.81E-02 | 5 | 2.51E-04 |
| **Top-HF∪Top-MD∪Top-Drug** | **0** | 1.00E+00 | **3** | 3.70E-01 | **5** | 1.53E-02 | **5** | 1.77E-02 |

**Supplementary Table 9**: Best-classifier proteins found in the Top-HF ∪ Top-MD ∪ Top-Drug set.

| | **Uniprot ID** | **Gene symbol** | **Gene name** |
|---|---|---|---|

| | | | |
|---|---|---|---|
| **LHF-HHF+** | P28482 | MAPK1 | Mitogen-activated protein kinase 1 |
| | P27361 | MAPK3 | Mitogen-activated protein kinase 3 |
| | P02751 | FN1 | Fibronectin |
| **LMD+HMD-** | P18084 | ITGB5 | Integrin beta-5 |
| | O75787 | ATP6AP2 | V-ATPase M8.9 subunit |
| | Q02297 | NRG1 | Pro-neuregulin-1, membrane-bound isoform |
| | P06748 | NPM1 | Nucleophosmin |
| | P01583 | IL1A | Interleukin-1 alpha |
| **LMD-HMD+** | P04085 | PDGFA | Platelet-derived growth factor subunit A |
| | P02675 | FGB | Fibrinogen beta chain |
| | P05155 | SERPING1 | Plasma protease C1 inhibitor |
| | P05230 | FGF1 | Fibroblast growth factor 1 |
| | P42574 | CASP3 | Caspase-3 subunit p12 |

**Supplementary Table 10**: Intersection of several set of proteins defined with GUILDify with the biomarkers obtained from the TPMS analysis. The p-values are calculated using a Fisher's exact test. The p-values above 0.05 are remarked in red.

| **Sets of proteins** | **LHF ∩ LMD+ HMD-** | **P-value** | **LHF ∩ LMD- HMD+** | **P-value** |
|---|---|---|---|---|
| Drug seeds | 0 | 1.00E+00 | 0 | 1.00E+00 |
| Top-Drug | 1 | 2.90E-01 | 0 | 1.00E+00 |
| HF seeds | 1 | 1.45E-01 | 1 | 1.28E-01 |
| Top-HF | **2** | 4.03E-02 | 1 | 2.48E-01 |
| MD seeds | **2** | 1.80E-02 | **4** | 2.54E-05 |
| Top-MD | **4** | 2.75E-04 | **4** | 1.56E-04 |
| Top-HF∪Top-MD∪Top-Drug | **5** | 1.37E-03 | **5** | 6.89E-04 |

**Supplementary Table 11**: Biomarkers from the TPMS analysis found in the Top-HF ∪ Top-MD ∪ Top-Drug set.

| | **Uniprot ID** | **Gene symbol** | **Gene name** |
|---|---|---|---|
| **LHF ∩ LMD+ HMD-** | Q02297 | NRG1 | Pro-neuregulin-1, membrane-bound isoform |
| | P06748 | NPM1 | Nucleophosmin |
| | P01583 | IL1A | Interleukin-1 alpha |

| | | | |
|---|---|---|---|
| | P61981 | YWHAG | 14-3-3 protein gamma, N-terminally processed |
| | P18084 | ITGB5 | Integrin beta-5 |
| **LHF ∩ LMD-HMD+** | P05121 | SERPINE1 | Plasminogen activator inhibitor 1 |
| | P02675 | FGB | Fibrinogen beta chain |
| | P05230 | FGF1 | Fibroblast growth factor 1 |
| | Q15109 | AGER | Advanced glycosylation end product-specific receptor |
| | P05155 | SERPING1 | Plasma protease C1 inhibitor |

24. Burnett M. Blocking Brute Force Attacks. UVA Comput Sci.

25. Keinosuke Fukunaga. Introduction to statistical pattern recognition 2nd edition. 1990 . Epub ahead of print 1990. DOI: 10.1016/0098-3004(96)00017-9.

26. Haykin S. Neural networks: a comprehensive foundation. 1994 . Epub ahead of print 1994. DOI: 10.1017/S0269888998214044.

27. Madsen H, Thyregod P. A Generalized linear Model with binomial distribution and probit link function has been used as classifier. In: Introduction to General and Generalized Linear Models. Chapman & Hall/CRC. 2011.

28. Gorban AN, Zinovyev AY. Principal Graphs and Manifolds. In: Handbook of Research on Machine Learning Applications and Trends: Algorithms, Methods and Techniques. Information Science Reference. 2009; pp. 28–59.

29. Shimizu K, Short DA, Kedem B. Single- and Double-Threshold Methods for Estimating the Variance of Area Rain Rate. J Meteorol;71.

30. Kohavi R. A study of cross-validation and bootstrap for accuracy estimation and model selection. Proc 14th Int Jt Conf Artif Intell - Vol 2 1995;2:1137–1143.

31. BIPM. Guides in metrology, Guide to the Expression of Uncertainty in Measurement (GUM) and International Vocabulary of Metrology (VIM). 2008.
